# A framework for *in situ* molecular characterization of coral holobionts using nanopore sequencing

**DOI:** 10.1101/2020.05.25.071951

**Authors:** Quentin Carradec, Julie Poulain, Emilie Boissin, Benjamin CC Hume, Christian R Voolstra, Maren Ziegler, Stefan Engelen, Corinne Cruaud, Serge Planes, Patrick Wincker

## Abstract

Molecular characterization of the coral host and the microbial assemblages associated with it (referred to as the coral holobiont) is currently undertaken via marker gene sequencing. This requires bulky instruments and controlled laboratory conditions which are impractical for environmental experiments in remote areas. Recent advances in sequencing technologies now permit rapid sequencing in the field; however, development of specific protocols and pipelines for the effective processing of complex microbial systems are currently lacking. Here, we used a combination of 3 marker genes targeting the coral animal host, its symbiotic alga, and the associated bacterial microbiome to characterize 60 coral colonies collected and processed *in situ*, during the *Tara* Pacific expedition. We used Oxford Nanopore Technologies to sequence marker gene amplicons and developed bioinformatics pipelines to analyze nanopore reads on a laptop, obtaining results in less than 24 hours. Reef scale network analysis of coral-associated bacteria reveals broadly distributed taxa, as well as host-specific associations. Protocols and tools used in this work may be applicable for rapid coral holobiont surveys, immediate adaptation of sampling strategy in the field, and to make informed and timely decisions in the context of the current challenges affecting coral reefs worldwide.

## Introduction

Coral reefs are threatened worldwide by global environmental changes and local anthropogenic pressures [1]. Studying coral reef ecosystems today is essential to understand what their stressors are and how their biodiversity will be affected in the coming years. Coral holobionts are ecological units organized from an anthozoan cnidarian animal host (the coral) and its obligate photosynthetic dinoflagellate endosymbionts of the family Symbiodiniaceae [2]. These dinoflagellates are subject of numerous scientific studies given their predominant role in coral sensitivity and resilience to bleaching events [3]. However, an array of other organisms (Bacteria, Archaea, Fungi, Protists, and viruses) living within or around the coral colony may be as important as the dinoflagellate for coral holobiont health [4, 5]. For example, symbiotic cyanobacteria have been shown to be a source of nitrogen for the scleractinian coral *Montipora sp.* [6]. Conversely, several strains of *Vibrio* are putative causative agents of coral bleaching [7]. Development of high-throughput sequencing techniques have been fundamental in enabling our characterization of the taxonomic diversity that makes up the coral holobiont.

Morphological identification of coral species is challenging for the non-specialist and even for coral researchers, microscopy equipment is often required to distinguish between closely related species [8]. For the past 2 decades, genetic markers are used to help the identification of coral species or establish population structures of these intricate taxa. However, due to a slow nucleotide substitution rates for mitochondrial genes within anthozoans [9], common genetic markers such as the cytochrome c oxidase subunit I or the cytochrome b are not discriminant enough to identify corals at the species level [10, 11]. Nuclear markers like the 18S or the 5.8S rRNA [12, 13] are generally more variable between species although multi-locus and microsatellite analysis are often required to precisely determine species boundaries [14].

The most commonly used marker gene to study the diversity of coral symbionts in the Symbiodiniaceae diversity is the Internal Transcribed Spacer 2 (ITS2) region of the rDNA [15]. Whereas only 22 Symbiodiniaceae species are formally described, 432 different ITS2 sequences recently grouped in seven genera are today referenced and may encompass a much larger number of species [2]. ITS2 is a multi-copy marker that can resolve Symbiodiniaceae at the level of taxa and strains, but the high intragenomic diversity of the ITS2 sequence poses analytical challenges in distinguishing intra-from inter-genomic diversity [16]. Different strategies have been developed to solve this problem including a novel analytical framework (“SymPortal”). This tool makes explicit use of the intragenomic diversity by employing the resolution of next-generation-sequencing approaches to determine ITS2 type profiles of putative Symbiodiniaceae taxa based on consistent co-occurrence of defining intragenomic ITS2 variants [17]. The large diversity of Symbiodiniaceae symbionts and numerous different microhabitats across coral reefs make the global comprehension of environmental and biological drivers of host-symbiont specificity challenging [18-20].

The 16S rRNA gene is the most commonly used marker to assess the bacterial diversity in coral holobionts [21]. Previous quantification of coral-associated bacterial assemblage richness have identified up to 100,000 distinct OTUs dominated by gamma- and alpha-proteobacteria [22]. Among them *Endozoicomonas* is probably the most abundant and widely distributed bacterial genus [23, 24]. For some coral species, the bacterial composition is highly variable according to coral reef site, environmental conditions, or seasons [25], whereas for other corals, bacterial compositions is rather fixed and less variable [26]. As such, comprehensive sampling efforts are needed to discover all possible associations and to identify taxa of putative functional importance [27].

The recent development of the MinIon portable sequencer by Oxford Nanopore Technologies (ONT) allows real time long read sequencing and is practicable in the field. Barcoding experiments have been realized in various environments for the molecular identification of rare species [28-30]. However, very few studies have realize metabarcoding experiments on complex samples due to the lack of analysis pipelines able to work with long sequences and to manage the high error rate of nanopore sequencing for the taxonomic identification [31].

Here, we investigate the diversity of microbial assemblages associated with corals sampled around an isolated reef in north eastern Papua New Guinea (Kimbe Bay). Without access to bulky sequencing instruments and with limited laboratory facilities, we used the MinION device onboard the research vessel *Tara* to evaluate coral host identity as well as Symbiodiniaceae and bacterial composition of sampled coral specimens in the field. We used simultaneously three rRNA marker genes to characterize the coral holobiont: the full-length 18S rRNA for the coral host, the ITS2 for Symbiodiniaceae diversity, and the 16S rRNA for the bacterial assemblages. In total, 55 samples of scleractinian corals from 13 different genera and 5 samples of the fire coral *Millepora* (Hydrozoa) were assessed.

## Results

### Coral sample collection and nanopore sequencing of marker genes

A total of 60 coral colonies were collected at 4 sites in Kimbe bay, Papua New Guinea during a 12-day research trip in December 2017 onboard the *Tara* vessel as part of the *Tara* Pacific expedition (Figure 1). In order to identify Symbiodiniaceae and the bacterial community associated with each coral, we extracted DNA from each colony and sequenced 3 different marker genes on the Oxford Nanopore MinION device (Figure 2 and Methods). For each colony, the full-length 18S rRNA sequence (1.8 kb) was PCR-amplified with primers designed for coral identification [32], the ITS2 region (250bp) was amplified with primers specific for the Symbiodiniaceae family [16], and the full-length 16S rRNA (1.3 kb) with bacteria-specific primers [33]. 12 unique identifier sequences (barcodes) were added to the 5’ends of each primer pair. This strategy allowed us to sequence the holobiont of 12 coral colonies in a single MinION run lasting 24h (Figure 2).

**Figure 1:**
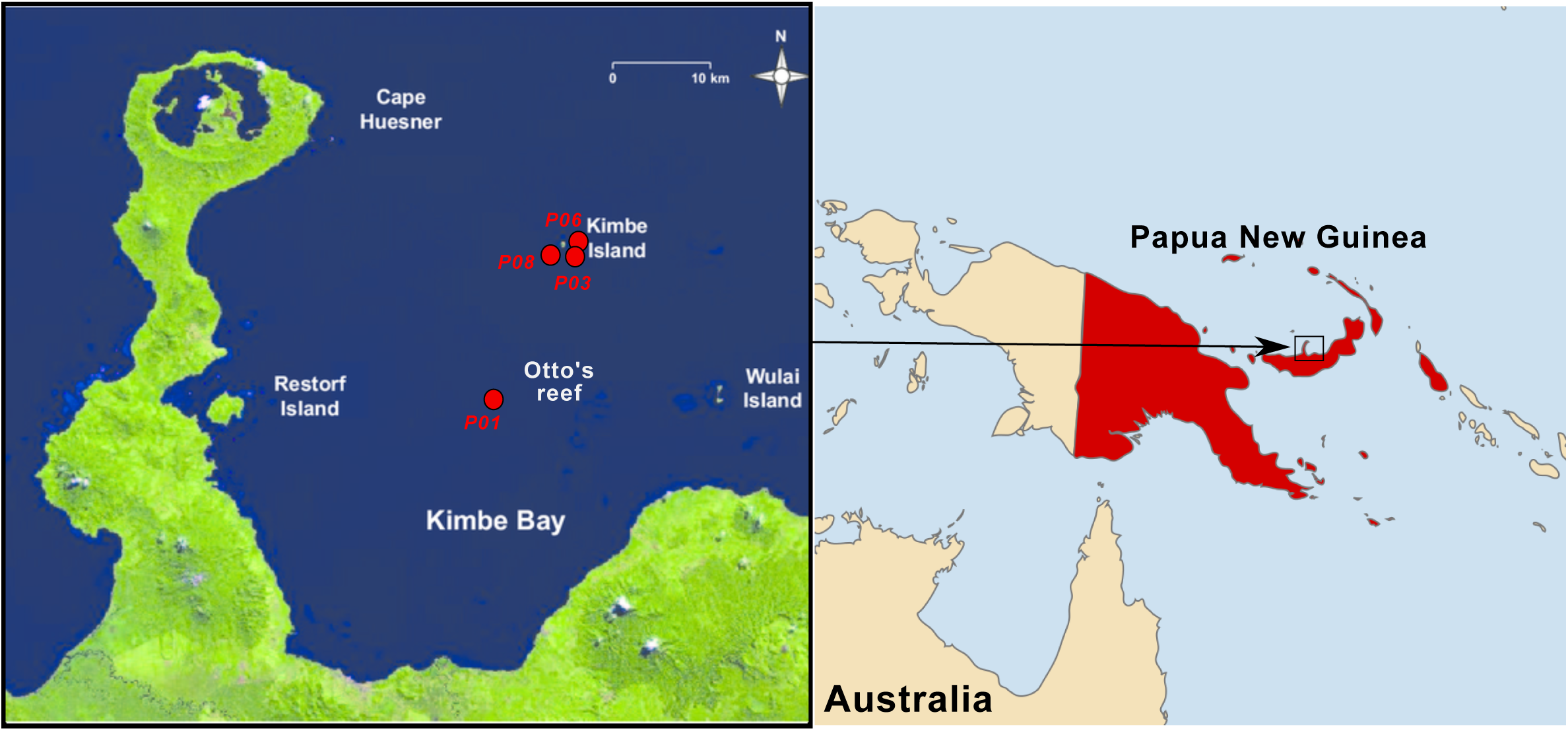
Map of the sampling sites. The four sampling sites are indicated by red dots on the Kimbe Bay map. The first site (P01) is close to Otto’s Reef, the three other sites (P03, P06 and P08) are around the Kimbe Island.

**Figure 2:**
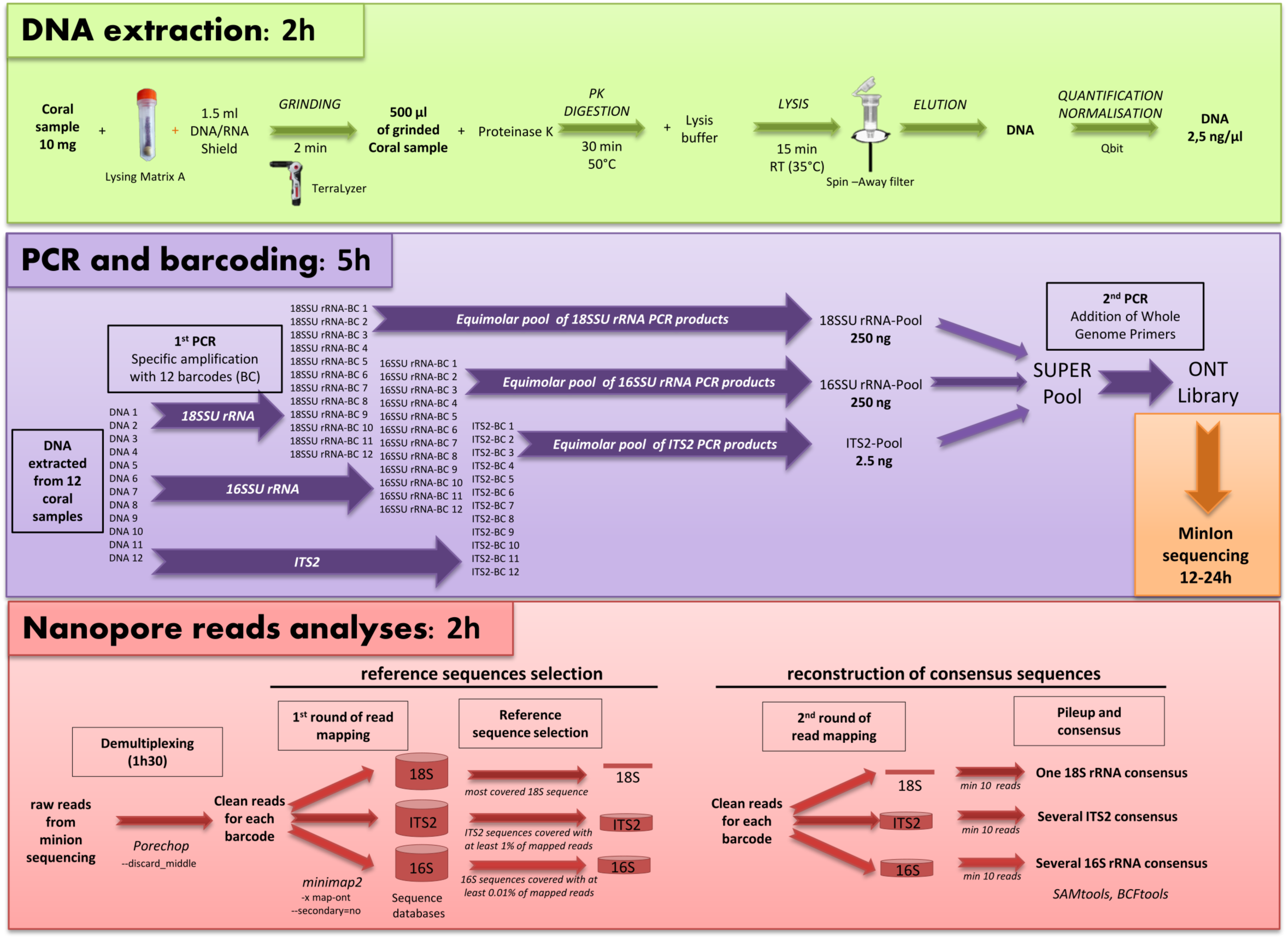
Pipeline for *in situ* analysis of coral holobionts. The approximate times for each step correspond to the processing of 12 coral samples. Protocols and tools parameters are detailed in the method section.

After 5 sequencing runs over a period of 10 days, we obtained a total of 2,019,607 reads (2.0 Gb) from 60 coral colonies and the unique barcodes were recognized for 32% of the sequences on average (Supplementary Table S1). For 48 samples, more than one thousand demultiplexed sequences were obtained for each barcode, which is seemingly sufficient to identify the coral as well as dominant Symbiodiniaceae and bacterial taxa. For the 12 remaining samples (11 sequenced during the last run), the number of reads was too low to identify the coral and the bacterial community. In this last run, most of the nanopore channels became inactive within a few seconds of run resulting in a very low output. This phenomenon is most probably due to the conservation of the flow cell for more than 2 months before the run which is longer than ONT specifications. However, the number of sequencing reads obtained was sufficient to study the ITS2 amplicon, we then decided to keep this MinION run for downstream analysis. The identification of 18S rRNA, ITS2 and 16S rRNA and sequences was realized by the mapping of nanopore reads against specific databases for each marker gene (see Methods).

### Identification of corals, Symbiodiniaceae, and bacterial communities

Corals were identified by both their morphological traits (Supplementary Figure S1) and the analysis of 18S rRNA sequences. A full-length 18S rRNA consensus was obtained for 50 corals covering 10 scleratinia families and the fire coral *Millepora*. The coral identification with the 18S rRNA was limited at the genus level for non-acroporid corals because the 18S rRNA sequence is not discriminant at species level for these genera [12] (Supplementary Table S2). A phylogenetic tree was reconstructed confirming the taxonomic identification of these corals (Supplementary Figure S2). The DNA extraction and/or the PCR amplification failed for 10 coral colonies. In these cases we based our identification on the morphological traits only.

In order to identify Symbiodiniaceae diversity in each coral colony, nanopore reads were mapped against a database of 432 *Symbiodinium* ITS2 sequences [18]. Several Symbiodiniaceae species may coexist in a coral colony so we can expect multiple ITS2 sequences per sample. So for each sample, ITS2 sequences covered with at least 1% of all nanopore reads aligned were conserved. In 53 of the sampled coral colonies we succeeded in identifying at least 1 ITS2 sequence. A similar method was used to characterize the bacterial community from the full-length 16S rRNA (see Methods).

In order to get a broad overview of the coral holobionts analyzed in this study, we constructed a force-directed graph (Figure 3). This network showed that Symbiodiniaceae taxonomy correlates to coral host at the family level, whereas bacterial specificity to its host is dependent on the bacterial taxa. Some bacterial taxa are detected in almost all coral hosts sampled (in the centre of Figure 3), while others appear host-specific.

**Figure 3:**
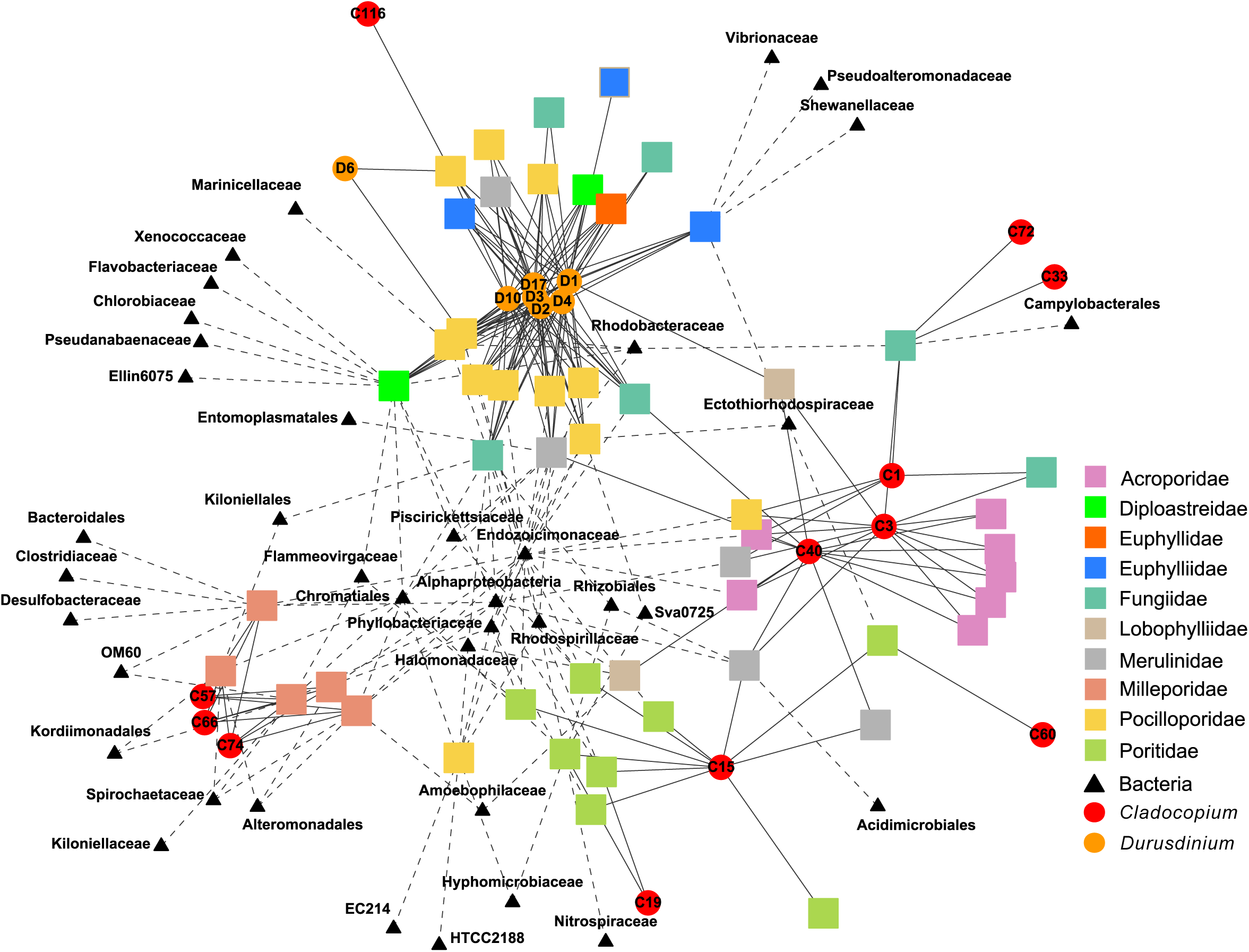
Coral holobiont network. Network representation of a force directed graph of Symbiodiniaceae and bacterial families living within or around each coral colony. Coral colonies are represented by a square and coloured according to their taxonomy (family level). Each node connected to a coral colony represents an organism living within or around the colony. Symbiodiniaceae taxa identified with nanopore sequencing are represented by a circle coloured by taxonomic origin; red for Cladocopium (clade C) and orange for *Durusdinium* (clade D). ITS2 sequences are indicated inside each circle. Each bacterial sequence is represented by a triangle and its family of origin is indicated below. Symbiodiniaceae ITS2 sequences and Bacteria 16S rRNA sequences covered with less than 5% and 1% of all mapped reads respectively are not represented in this figure.

### Symbiodiniaceae diversity in coral holobionts

Symbiodiniaceae ITS2 sequences from taxa in the genus *Durusdinium* (clade D) were dominant in 22 coral colonies including 4 Fungiidae colonies and all *Diploastrea, Galaxea*, and *Pocillopora* (Figure 3 and Figure 4a). A total of 8 different ITS2 sequences were reconstructed from these samples. The D1 (*D. glynni*) is the relatively most abundant sequence representing between 40% and 60% of reads in each coral colony. D2, D3, D4, D10, and D17 were recovered in all of these colonies in lower proportions suggesting non-specific alignment of ONT reads. To test this hypothesis, we re-sequenced the same PCR amplicons using Illumina technologies, which have a reduced sequencing error rate (Figure 4b). Sequencing results confirmed the dominance of the D1 sequence in these 22 coral colonies, but also revealed distinct patterns of less abundant, i.e. minor, ITS2 sequences: D4 in Fungiidae, *Diploastrea* and *Galaxea* colonies, and D2 in *Pocillopora* colonies. These differences between ONT and Illumina reveal a limitation for the identification of minor ITS2 sequences of the genus *Durusdinium* with the current error rate of ONT sequences.

**Figure 4:**
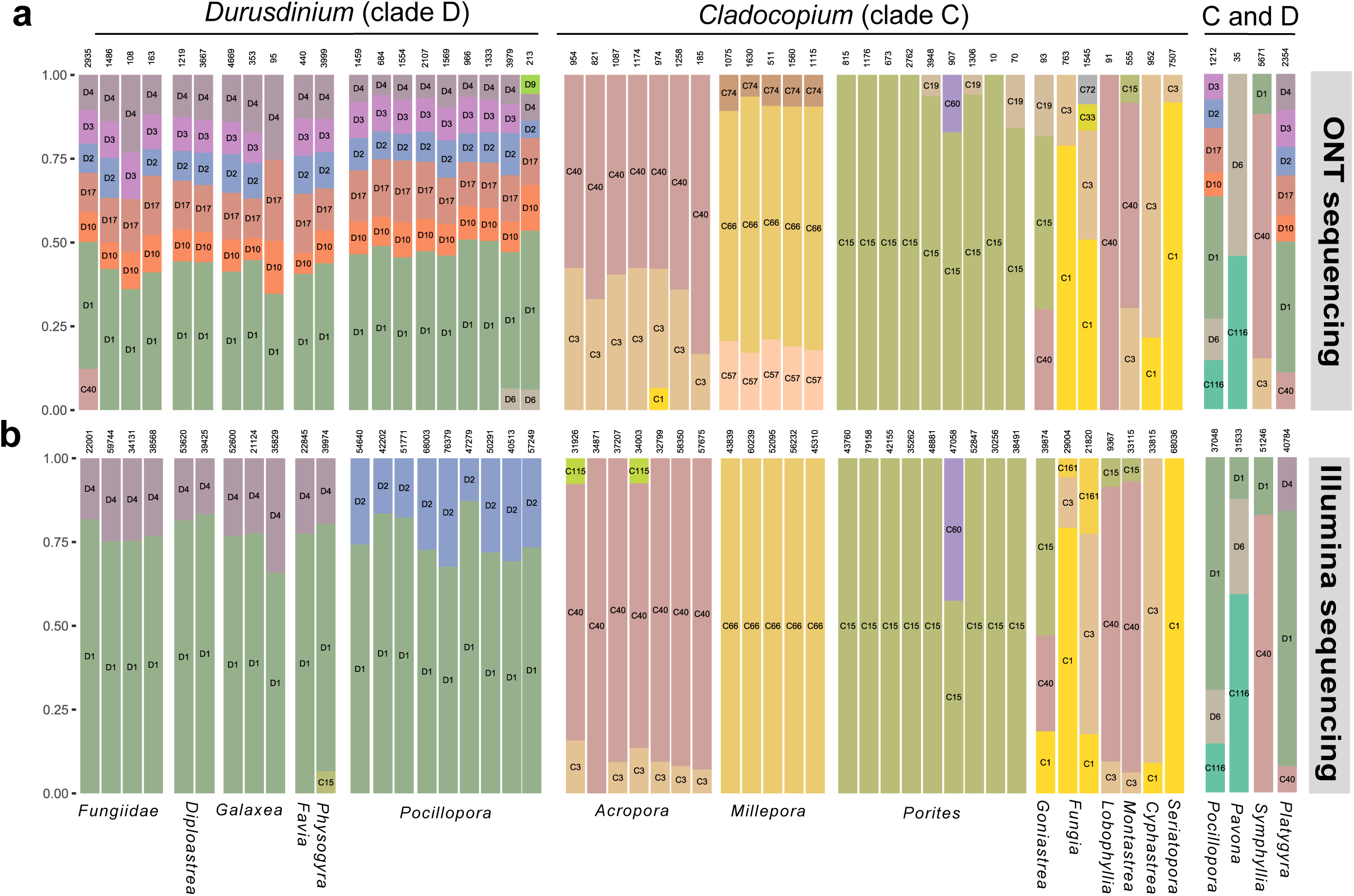
Symbiodiniaceae ITS2 diversity in coral colonies sequenced with ONT or Illumina technology. Each coloured bar represents the proportion of Symbiodiniaceae ITS2 detected in a coral colony which have a relative abundance above 5%. The total number of reads is indicated on top of each bar. The name of each ITS2 sequence is indicated and represented by a specific color. Corals colonized by Symbiodiniaceae of *Cladocopium* genus are on the left panel, *Durusdinium* genus on the middle panel and by both genera on the right panel. a) PCR amplicons sequenced with ONT. b) Same PCR amplicons sequenced with Illumina technology.

Several sequences representing *Cladocopium* taxa (clade C) were detected in different coral families (Figure 3 and Figure 4a). The C3 and C40 sequences (1 insertion and 1 substitution between them) were present in all *Acropora* colonies. Their co-presence is confirmed with the Illumina sequencing (Figure 4b and Supplementary Figure S3). Three *Cladocopium* sequences were also recovered from the *Millepora* colonies: C57, C66, and C74 in consistent proportion (9% C74, 72%, C66, 19% C57) (Figure 4a). C57 and C74 variants (respectively 1 and 2 substitutions with C66) were not observed with Illumina sequencing suggesting that their detection is also due to the error rate of ONT sequences. We detected C15 sequences in all *Porites* colonies and C60, a C15-derived sequence (2 substitutions) confirmed by Illumina sequencing, in 1 colony (Figure 4a). Finally, 7 coral colonies from 6 different genera present a large diversity of *Cladocopium* ITS2 variants, however several samples of the same coral species would have been necessary to substantiate these associations (Figure 4a).

Sequences from the *Cladocopium* and *Durusdinium* genera were found co-occurring in 4 colonies: one *Plathygyra* with the C40 and D1, one *Symphyllia* colony with C3, C40 and D1, one *Pavona* colony with the C116 and the D6 and one *Pocillopora* with C116 and D1 ITS2 sequences (Figure 4). The co-occurrence of these two genera in a coral colony were previously reported in Papua New Guinea [34].

### Coral-associated bacterial assemblages

We retrieved 1,637 unique bacterial 16S sequences belonging to 77 orders of bacteria. Among these sequences, 175 are detected in 2 or more coral colonies. These Bacteria were identified in 31 coral colonies (Supplementary Table S2 and S3). Of the coral colonies sampled, *Oceanospirillales* was the most common bacterial order (detected in 22 coral colonies), followed by *Rhizobiales* (12), uncharacterized alphaproteobacteria (12) and *Chromatiales* (11) (Supplementary Table S2 and S3). These marine bacteria are commonly detected in a large diversity of corals colonies and are known to be associated with coral tissue and mucus. For instance, *Endozoicomonadaceae* (*Oceanospirillales*) are known to be abundant in healthy mucus and absent or in very low abundance in diseased corals [25, 35-38]. Their presence in a large number of corals (20 colonies including *Poritidae, Pocilloporidae, Acroporidae*, and *Milleporidae*) may indicate the health of this coral reef, although the functional role of *Endozoicomonas* remains elusive [23, 39]. Among the 186 unique full-length 16S sequences belonging to the *Endozoicomonas* family, several are shared between different coral hosts (maximum of 12 samples) showing that the family is large, commonly found in corals, but also display a pattern of fine-scale genetic differentiation with host (Supplementary Table S3) as recently shown [40, 41]. In addition, *Ectothiorhodospiraceae* family and *Kordiimonadales* order detected in several samples were so far not commonly identified within corals. From our data, we argue that they may represent important families, either for this particular reef or in a broader context, awaiting further studies.

At the same time, we found opportunistic bacterial taxa known to be associated with corals under stress. For instance, a bacterium belonging to the *Vibrionaceae* family was detected in one colony of *Galaxea* (*Euphylliidae*). This family of bacteria has been described to be prevalent in diseased corals [42, 43]. Furthermore, two *Alteromonadales* species were detected in this colony. *Shewanellaceae*, already described in *Favia* corals [44], and *Pseudoalteromonadaceae* describe in corals affected by sedimentation and local sewage [45] suggests that this colony may be under stress.

An endosymbiotic bacterium of *Millepora* belonging to *Spirochaetaceae* family was detected in four out of five coral colonies and a *Kordiimonadales* bacterium in two colonies, these bacterial families were already described in healthy as well as sick tissues of *Millepora* [46, 47]. The Gammaproteobacteria *Congregibacter* (OM60, *Alteromonadales*) was also detected in two colonies; this photosynthetic bacterium is found abundant in coastal ecosystems, but was never reported in association with *Millepora* [48]. Three other bacterial families (*Clostridiaceae, Desulfobacteraceae* and *Bacteroidales*) were detected in one sample of *Millepora* (P03-C051) suggesting a diseased colony (Figure 2 and Supplementary Table S2) [49, 50]. The aspect of this *Millepora* colony with the presence of parasites, the strong space competition with other coral species and the presence of dead parts colonized by algae corroborate this hypothesis based on the bacterial composition (Supplementary Figure S1).

## Discussion

Recent studies have shown the efficiency of Nanopore sequencing technology for rapid species identification in the field in remote locations [28, 29]. However, this technology has so far not been applied to characterize complex ecosystems or holobionts. In this study, the MinION device was used to describe the diversity of one of the most diverse ecological units on earth: the coral holobiont [51].

A discriminant marker gene for all corals is still lacking, but the full-length 18S rRNA sequence used in this study was sufficient to identify corals at the species level for acroporids and at the genus level for most of the other corals. Coral identification with 18S rRNA sequencing in the field may open coral studies to non-specialist in contrary to the morphological identification that requires taxonomic expertise. Moreover, the identification in less than 24h on MinION device could significantly evolve sampling methods for corals. Given the ongoing uncertainties with coral taxonomy designations at the species level, it should be noted that even a designation of corals to the genus level, may be considered a big step forward with regard to diversity assessment to aid conservation efforts.

In addition to the identification of the coral host species, we successfully characterized the Symbiodiniaceae community and recovered sequences from the dinoflagellate genera *Cladocopium* and *Durusdinium. Cladocopium* symbiont genus is the most diverse of the Symbiodiniaceae family in the Arabian Seas, the Indo-Pacific, and the Atlantic-Caribbean [20, 52, 53]. This diversity is mainly driven by rapid host specialization, but also by specific environmental conditions [54], suggesting that the diversity may be best explained by local adaptation to the environmental condition of this remote reef. If correct, this suggests sampling more coral reefs is essential to get a complete view of all possible Symbiodiniaceae symbionts for a coral species. Sequences belonging to the *Symbiodinium* genus (clade A) were not detected in any coral colony sampled in Kimbe Bay, corroborating previous observations in PNG [34]. The comparison between the two sequencing technologies has reveal a limitation in the identification of the symbiont taxa with ONT. Although the dominant symbiont in each coral has always been correctly assigned in our study, rare taxa are sometimes mis-assigned due to the low sequencing depth and the high error rate of nanopore sequencing.

Regarding the coral-associated bacterial assemblages, we observed highly distinct phyla. Among those, we found common marine bacteria associated to coral mucus as well as more specific endosymbiotic relationships. Although the sequencing depth is insufficient to detect rare bacterial taxa, our results support that bacterial community composition assessed with the current technology may be used as an indicator of coral health.

The small size of the MinION sequencer coupled to a simple laptop is particularly convenient for use on a research vessel where the work space and electricity use are extremely limited. At present, this device is the only one able to execute a complete sequencing run under these conditions. A −20°C freezer is sufficient to conserve reagents for DNA extraction, ONT library preparation, and MinION sequencing, allowing molecular experimentations during long-term expeditions in distant islands in total autonomy. In addition, several new developments ongoing by ONT to lyophilize sequencing reagents will be useful improvements to enable room temperature storage of reagents for several months.

The bioinformatic pipeline developed in this article can be run on a simple laptop without internet access, so long as reference databases are prepared and downloaded before the expedition. The method described in this study will significantly improve the capability of local surveys and enable researchers in remote locations to make informed and effective sampling and experimental decisions while in the field. This approach also carries an element of capacity building in remote areas, as it allows local users to have full access over the data generated and analysed. In remote places, this method could be performed to rapidly evaluate the state of numerous coral colonies on a reef in terms of holobiont diversity and then orient further sampling in the most interesting features like the presence of unexpected bacteria or Symbiodiniaceae or rare coral species.

In addition, ONT allows the sequencing of full marker genes instead of short amplicon regions, often insufficient for resolving taxonomies to the species level. Long-read sequencing have already shown their efficiency on coral holobiont characterization [33]. The main limitation of this technology is the current error rate that we aimed to address here through the generation of consensus sequences from reads aligned on each sequence of the database. Despite this analysis pipeline, very similar sequences diverging by a single nucleotide could be assigned incorrectly as was the case for minor variants of *Durusdinium* D1 and *Cladocopium* C66. Future development could use the Intramolecular-ligated Nanopore Consensus (INC-Seq) method to correct remaining sequencing mistakes and allow *de novo* identification of sequences without reliance on reference databases [55]. The expected improvements in error rates will impact the bacterial detection by diminishing the number of reads necessary to confidently build a consensus, which is presently high. This would lead to delineation of a more complex microbiome.

Coral holobionts require large sampling efforts to obtain an accurate and complete network of the diversity of coral-associated Symbiodiniaceae and bacteria present in a reef. The here-presented study realized in Papua New Guinea demonstrates the efficacy of the nanopore method for a rapid survey of a large number of coral colonies which could help future evaluation of threatened coral reefs and holds applications for monitoring with a substantial component of capacity building.

## Materials and Methods

### Coral and Hydrozoa sampling and DNA extraction

60 coral samples were collected between 5 m to 20 m depth by removing a fragment on four different reefs of the Kimbe Bay in Papua New Guinea (New Britain island) (Figure 1). Various families of scleractinian corals as well as the hydrozoan genus *Millepora* were collected (Supplementary Figure S1). Coral fragments were placed in zip-lock bags under water then stored in 2 ml RNAse/DNAse free tubes containing Lysing Matrix A (MP Biomedical) and 1.5 ml of DNA/RNA shield preservative buffer (Zymo Research, Irvine, California, USA). Samples were then placed in a Terralyzer Instrument (Zymo Research) and grinded during 2 minutes. Grinded samples (500 µl) were incubated 30 minutes at 55°C with 75 µl of proteinase K. Then, 1.5 ml of Lysis Buffer (ZR-DuetDNA/RNA Miniprep Plus, Zymo Research) was added for 15 minutes at room temperature (around 35°C). Coral DNA was then extracted using the ZR-DuetDNA/RNA Miniprep Plus Kit following the manufacturer’s instructions. DNA was quantified on a Qubit dsDNA HS Assays (Thermo Fisher Scientific, Waltham, USA). High resolution coral pictures were archived at the European Bioinformatics Institute.

### Full-length marker gene amplification

The ITS2, 18S and 16S rRNA sequences were targeted for amplicon generation. For each primer, the ONT tail and one unique barcode (out of twelve) were added. Full-length 18S rRNA primers: TTTCTGTTGGTGCTGATATTGC-Barcode-AACCTGGTTGATCC TGCCAGT for the forward and ACTTGCCTGTCGCTCTATCTTC-Barcode-TGATCCTTCTGCAGGTTCACCTAC for the reverse primer [32]. Bacterial-specific 16S rRNA primers: 27F (TTTCTGTTGGTGCTGATATTGC-Barcode-AGAGTTTGATCMTGG CTCAG) and 1492R (ACTTGCCTGTCGCTCTATCTTC -Barcode-TACGGYTACCTTGTTA CGACTT) [33]. SYM_VAR ITS2 primers: TTTCTGTTGGTGCTGATATTGC-Barcode-GAATTGCAGAACTCCGTGAACC for the forward and ACTTGCCTGTCGCTCTATCTTCT-Barcode-CGGGTTCWCTTGTYTGACTTCATGC for the reverse [16]. PCRs were performed on board using 25 ng of DNA from each coral sample with the Advantage 2 kit (Takara Clontech) and a final primer concentration of 0.5 μM in a final reaction volume of 50 μl. PCR conditions were optimized to be able to generate the three amplicons using the same thermocycling program: initial denaturing at 95 °C for 1 min, 30 cycles each at 95 °C for 30 s, 55°C for 30 s, and 68 °C for 1 m, followed by a final extension step at 68 °C for 10 min. Twelve barcodes were available per primer set and allowed us to process DNAs from 12 samples in parallel on one 96 well PCR plate.

PCR products for each sample were pooled by targeted gene in equimolar ratios then each pool was 1:10 diluted and subsequently cleaned with 1 volume of AMPure XP (Beckman Coulter, Brea, California, USA) before quantification. 2.5 ng of ITS2-PCRs-pool, 250 ng of 16S-PCRs-pool and 250 ng of 18S-PCRs-pool were mixed to constitute the input of the ONT sequencing library. The smaller amount of ITS2-PCRs-pool was aimed at compensating for amplification bias, as PCR products for shorter sequences would result in a higher number of molecules sequenced.

### ONT library preparation and sequencing

Sequencing libraries were prepared for R7.9 flow cells run (FLO-MAP107) on MinION device using the Low Input by PCR Sequencing Kit SQK-LWP001 according to the four-primers PCR protocol from ONT with slight modifications detailed below. In order to add the Whole Genome Primers, a second multiplex PCR was performed from 500 ng of the PCR pool and a final primer concentration of 0.5 μM in a reaction volume of 50 μl. The following PCR conditions were used: initial denaturing at 95 °C for 3 min, 15 cycles at 95 °C for 30 s, 56°C for 30 s, and 68 °C for 1 m, followed by a final extension step at 68 °C for 10 min. PCR products were cleaned with 0.8 volume of Ampure XP and finally eluted in 20 µl of 10mM Tris-HCl pH8 with 50 mM NaCl. 50-100 fmol of PCR products were diluted in 9 µl then mixed with 1 µl of Rapid 1D Adapter and 1 µl of Ligase T4 Blunt (New England Biolabs, Ipswich, MA, USA). Pre-sequencing mix was incubated 10 min at 25°C and left on ice until ready to load. After priming the flow cell according to the manufacturer’s recommendations, 11 µl of pre-sequencing mix was combined with 30.5 μl of Running Buffer with Fuel Mix Buffer, 7 µl of water and 26.5 µl of Library Loading Beads and applied to the flow cell.

The sequencing run was performed for 12 to 24 hours. Bases were called during the MinION run with the MinKnow software (v. 1.7.14). The demultiplexing and adaptor trimming were done with porechop tool (https://github.com/rrwick/Porechop) with the option discard_middle. Sequences are archived at the European Bioinformatics Institute.

### Read mapping, consensus reconstruction, and species identification

Three specific databases were used to identify each set of nanopore reads. The coral 18S rRNA database contains full-length 18S rRNA for 140 scleratinia and 2 *Millepora [12]*. Five 18S rRNA sequences of Symbiodiniaceae (gi:176088, 12247076, 12247077, 148734588 and 12247080) were added in the database in order to detect and remove undesirable Symbiodiniaceae 18S amplified. The ITS2 database contains 432 sequences of Symbiodiniaceae [18] and finally the Greengenes database (v.13.5, http://greengenes.lbl.gov) was used to detect 16S rRNA reads. All nanopore reads were mapped on each database with minimap2 (v 2.0-r191) with the pre-set options “map-ont” [56]. For coral identification, the reference sequence that had the most sequences mapped to it was the only one retained in each sample. Then a second round of mapping (same parameters) was done on the selected reference in order to aggregate reads potentially mis-assigned during the first round of mapping. Two Maximum Likelihood phylogenetic trees with newly reconstructed sequences and the coral 18S rRNA database were further reconstructed, one with all sequences except Acroporidae and the other with Acroporidae only. For Symbiodiniaceae and the bacterial community the same strategy was conducted except that all references covered with a minimum of 1% (for ITS2 sequences) or 0.01% (for 16S rRNA sequences) of all reads mapped were kept for the second round of mapping. In addition for the bacterial community, reads mapped on eukaryote chloroplastic 16S were removed after the first mapping. SAMtools and BCFtools were used to reconstruct consensus sequences for each reference sequence covered with more than 10 nanopore reads with the following programs and options: mpileup -B -a -Q 0 –u; bcftools call -c --ploidy 1;vcfutils.pl vcf2fastq. All steps were proceeded on board of the research vessel *Tara*, at sea.

### Illumina sequencing and analysis of Symbiodiniaceae amplicons

The same PCR pools obtained onboard *Tara* were, later in laboratory, used for the sequencing of ITS2 amplicons on Illumina instruments in order to compare with ONT sequencing. 100 ng were directly end-repaired, A-tailed and ligated to Illumina adapters on a Biomek FX Lab Auto Workstation (Beckman Coulter). Then, library amplification was performed using Kapa Hifi HotStart NGS library Amplification kit and purified with AMPure XP (1 volume). Libraries concentrations were normalized to 10 nM by addition of Tris-Cl 10 mM, pH 8.5 and then applied to cluster generation according to the Illumina Cbot User Guide (Part # 15006165). Libraries were sequenced on a MiSeq instrument with paired end (2 × 300 bp). The taxonomic assignation of Illumina reads was done with the “SymPortal” pipeline [17]. In addition, novel ITS2 sequences were matched against the ITS2 database with BLAST (v2.6.0) then assigned to their closest match.

### Data availability

High resolution coral pictures and sequenced reads are accessible at the European Bioinformatics Institute repository under the BioProject PRJEB32905. All accession numbers are indicated in Supplementary Table S1.

## Acknowledgements

This project has been funded through the *Tara* Pacific consortium, France Genomique grant number ANR-10-INBS-09, and the Genoscope/CEA. We are keen to thank the commitment of the people and the following institutions and sponsors who made this singular expedition possible: CNRS, CSM, PSL, KAUST, Genoscope/CEA, ANR-CORALGENE, agnès b., the Veolia Environment Foundation, Region Bretagne, Serge Ferrari, Billerudkorsnas, AmerisourceBergen Company, Lorient Agglomération, Oceans by Disney, the Prince Albert II de Monaco Foundation, L′Oreal, Biotherm, France Collectivites, Kankyo Station, Fonds Francais pour l′Environnement Mondial (FFEM), Etienne Bourgois, UNESCO-IOC, the Tara Foundation teams and crew. Tara Pacific would not exist without the continuous support of the participating institutes.

## Author Contributions

JP, QC and PW designed the study. QC wrote the paper with substantial input from JP, BH, EB, CV, MZ, SP and PW. EB collected coral samples. JP coordinated *in situ* experiments and nanopore sequencing assisted by QC. QC, BH, MZ and EB performed barcoding data analyses. JP, QC, CC and SE contributed to the development of protocols and tools for nanopore sequencing and analysis. All authors discussed the results and commented on the manuscript.

## Additional Information

The authors declare no competing interests.

