## Supplementary Figure S1 for "A framework for *in situ* molecular characterization of coral holobionts using nanopore sequencing"

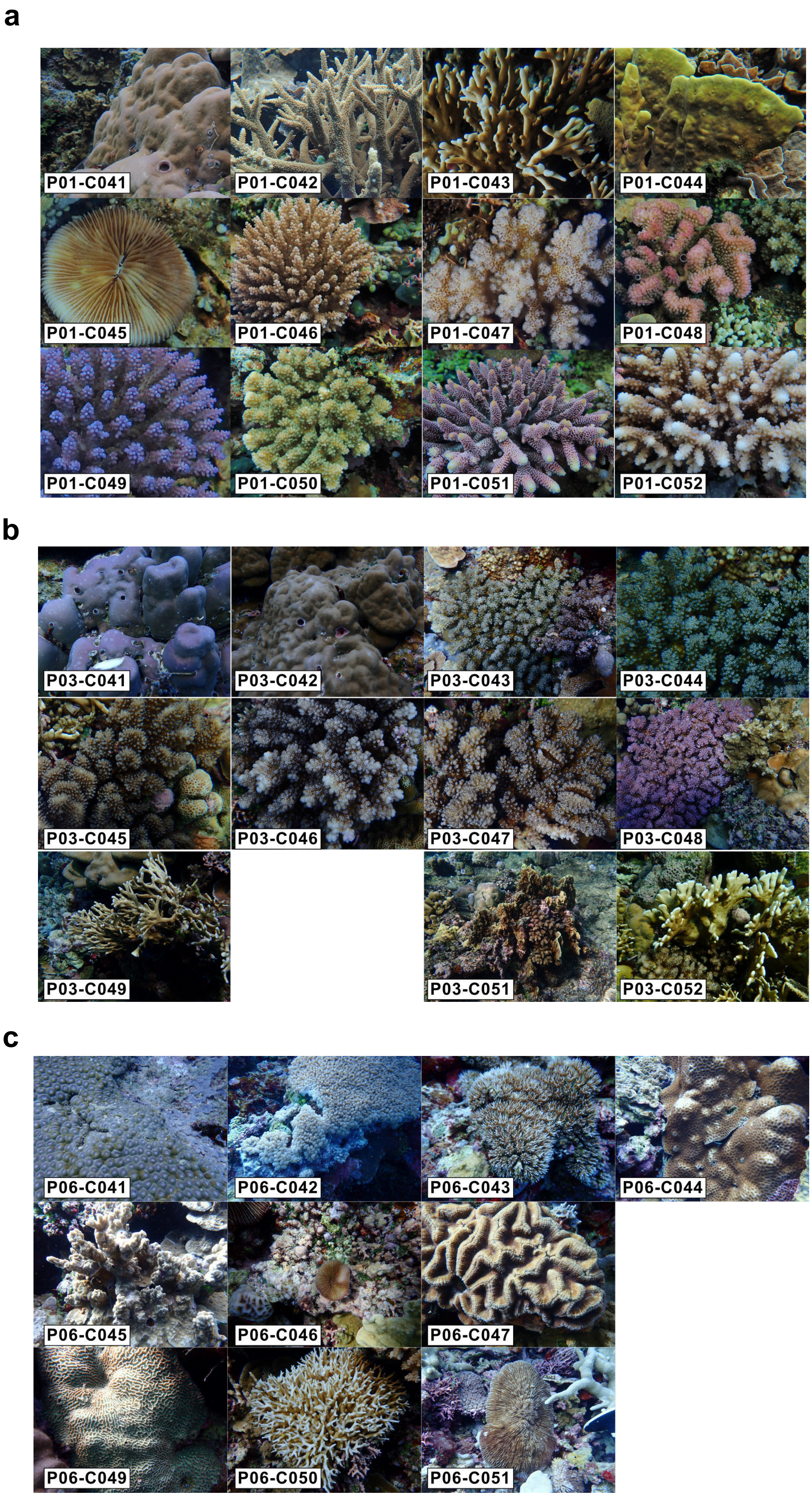
a
P01-C041
P01-C042
P01-C043
P01-C044
P01-C045
P01-C046
P01-C047
P01-C048
P01-C049
P01-C050
P01-C051
P01-C052
b
P03-C041
P03-C042
P03-C043
P03-C044
P03-C045
P03-C046
P03-C047
P03-C048
P03-C049
P03-C051
P03-C052
c
P06-C041
P06-C042
P06-C043
P06-C044
P06-C045
P06-C046
P06-C047
P06-C049
P06-C050
P06-C051

**d**

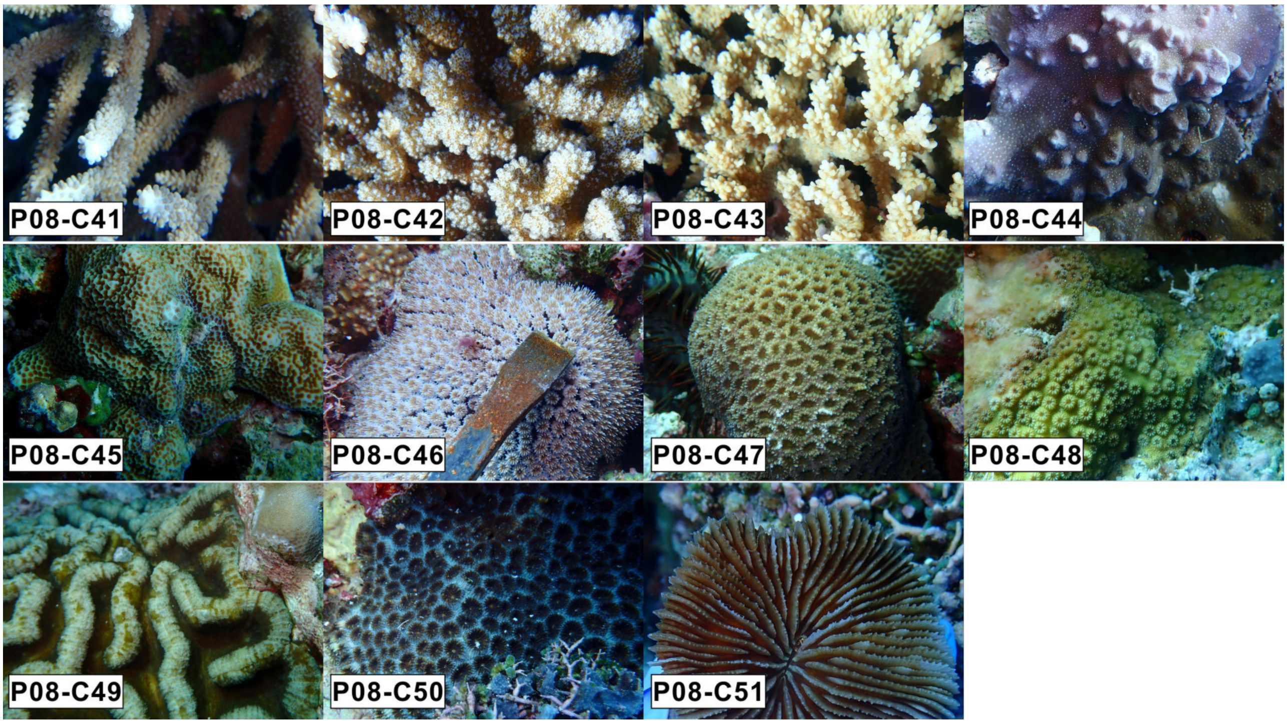


**e**

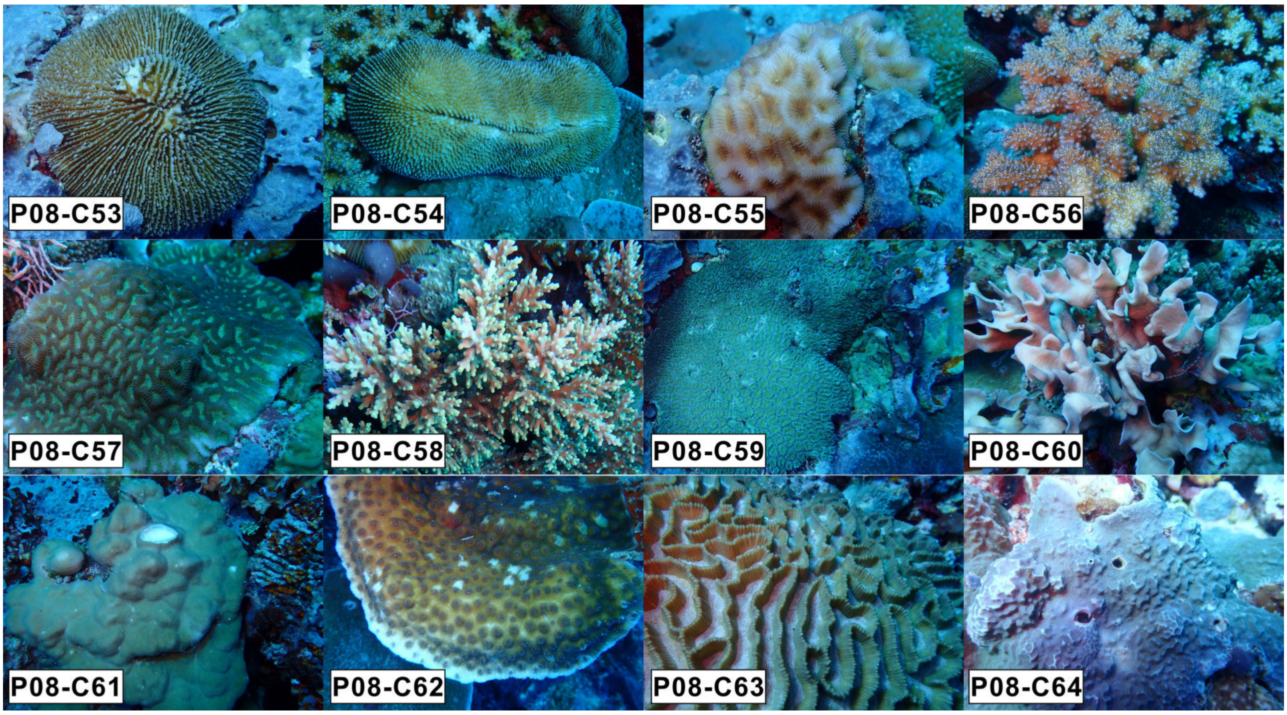


**Supplementary Figure S1:Pictures of sampled coral colonies. a)** P01-Run1. Porites lobata (C41), Acropora (C42), Millepora dichotoma (C43), Millepora platyphylla (C44), Fungia (C45), Pocillopora damicornis (C46), Acropora (C47), Pocillopora meandrina (C48), Pocillopora verrucosa (C49), Acropora (C50), Acropora (C51), Acropora (C52). **b)**. P03-Run2. Porites (C41), Porites lobata (C42), Pocillopora verrucosa (C43), Pocillopora verrucosa (C44), Pocillopora (C45), Pocillopora meandrina (C46), Pocillopora verrucosa (C47), Pocillopora (C48), Millepora dichotoma (C49), Millepora platyphylla (C51), Millepora dichotoma (C52). **c)** P06-Run3. Diploastrea (C41), Physogyra (C42), Galaxea (C43), Favites (C44), Porites rus (C45), Fungia (C46), Symphyllia (C47), Platygyra (C49), Seriatopora (C50), Fugiidae (C51). **d)** P08-Run4. Acropora (C41), Pocillopora verrucosa (C42), Acropora (C43), Porites rus (C44), Porites (C45), Galaxea (C46), Montastrea (C47), Cyphastrea (C48), Lobophyllia (C49), Favia (C50), Fungia (C51). **e)** P08-Run5. Fungia (C53), Fungia (C54), Pocillopora (C56), Goniastrea (C57), Acropora (C58), Galaxea (C59), Pavona cactus (C60), Porites Lobata (C61), Echinopora (C62), Oulophyllia (C63), Porites (C64).
