## Supplementary Figure S2 for "A framework for *in situ* molecular characterization of coral holobionts using nanopore sequencing"

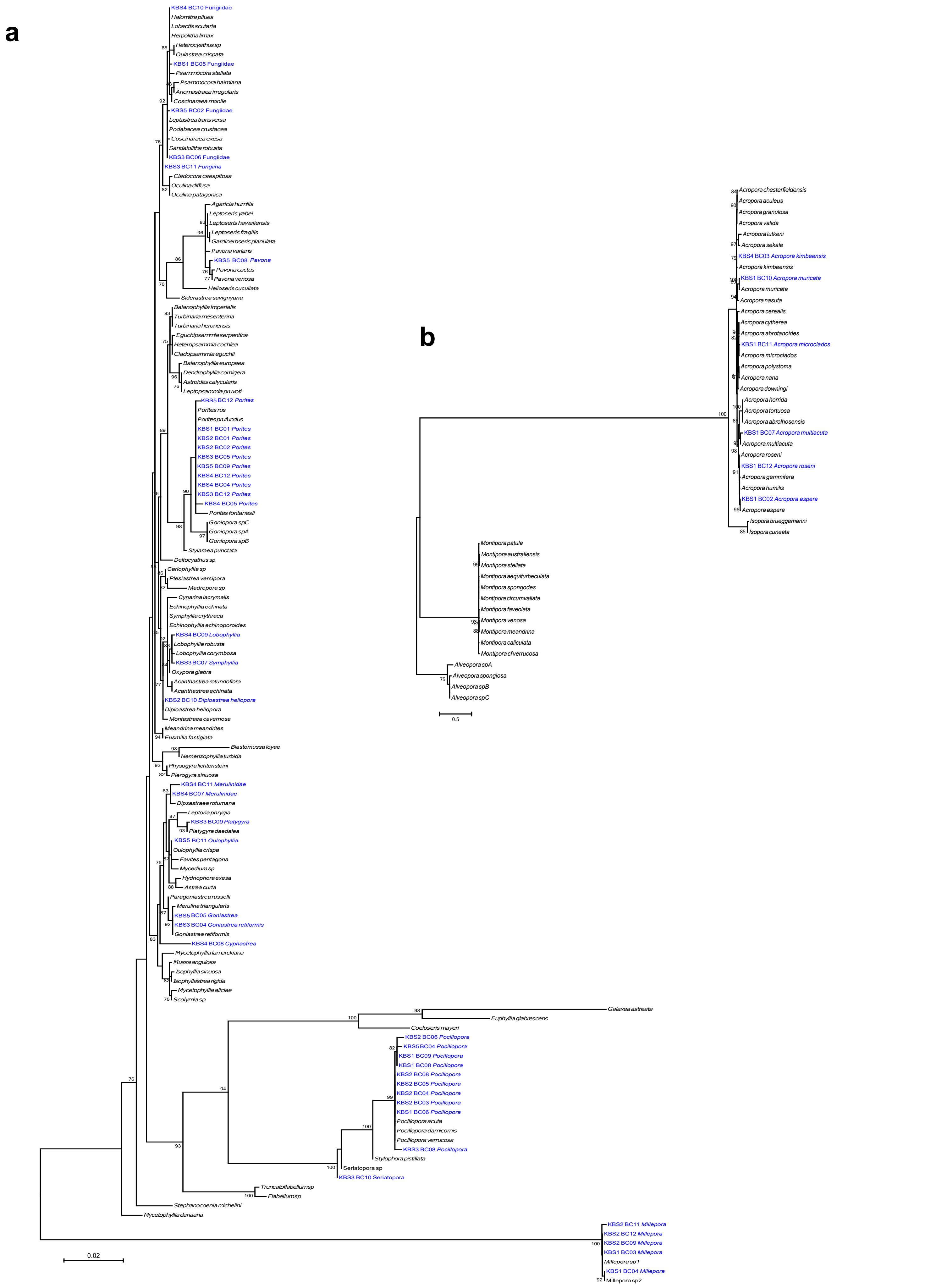


**Supplementary Figure S2:** Maximum Likelihood phylogenetic tree of coral 18S rRNA sequences. Sequences reconstructed in this study together with all 18S rRNA sequences from Arrigoni et al. 2018 are represented (12). (a) All corals except Acroporidae. (b) All Acroporidae 18S rRNA.
