## Supplementary Figure S3 for "A framework for *in situ* molecular characterization of coral holobionts using nanopore sequencing"

***Durusdinium* D1**

T(D10)

AACCAATGGCCCCC**T**GAA**C**G**C**GCATTGCACTCTTGGGACTTCCTGAGAGTATGT

G(D22)T(D17)

TTGCTTCAGTGCTTATTTTACCTCCTTGCAAGGTTCTGTCGCAACCTTGTGCCC

G(InsD6)

TGGCCAGCC**A**TGGGTTAACTTGCCCATGGCTTGCTGAGTAGTGATCTTTTAGAG

C(D6,D2,D1a)

CAAGCTCTGGCACGCTGTTGTTTGAGGCAGCCTATATTGAGGCTATTTCAAATG

ACGTTGCTACAAGCTTGATGTGTCCTTCTGCGCCGTTGCGCA**T**CCCATAGCATGA

A(D1.6)

***Cladocopium* C40**

AACCAATGGCCTCCTGAACGTGCGTTGCACTCTTGGGATTTCCTGAGAGTATGTCTGCTT

CAGTGCTTAACTTGCCC**C**AACTTTGCAAGCAGGATGTGTTTCTGCCTTGCGTTCTTATGA

*(C3b)

GCTATTGCCCTCTGAGCCAATGGCTTGTTAATTGCTTGGTTCTGGCAAAATGCTTTGCGC

G(InsC40b)

GCTGTTATTCAAGTTTCTACCTTCGTGGTTTTACTTGAGTGACGCTGCTCATGCTTGC**G**A

(C3b)A

CCGCTGGGATGCAGGTGCATGCCTCTAGCATGAAGTCA**G**ACAA

***Cladocopium* C66**

AACCAATGGCCTCCTGAACGTGCGTTGCACTCTTGGGATTTCCTGAGAGTATGTCTGCTT

**C**AGTGCTTAACTTGCCCCAACTTTGCAAGCAGGATGTTTTTCTGCCTTGCGTTCTTATGA

T(C74)

GCT**A**TTGTCCTCTG**C**GCCAATGGCTTGTTAATTGCTTGGTGCTTGCAAAATGCTTTGCGC

G(C57a) T(C57C57a)

GCTGTTATTCAAGTTTCTACCTTCGTGGTTTTACTTGAGTGA**C**GCTGCTCATGCTTGCAA

T(C74)

CCCGCTGGGATGCAGGTGCATGCCTCTAGCATGAAGTCAGACAA

***Cladocopium* C15**

AACCAATGGCCTCCTGAACGTGCGTTGCA**C**CCTTGGGATTTCCTGAGAGTATGTCTGCTT

T(C19) T(C2)

CAGTGCTTAACTTGC**C**CCAACTTTGCAAGCAGGATGTGTTTCTGCCTTGCGTTCTTATGA

*(C19)

GCTA**T**TGCCTTCTGC**G**CCAATGGCTTGTTAATTGCTTGGTTCTTGCAAAATGCTTTG**C**GC

C(C60) C(InsC15f) (C60)T

GCTGTTATTCAAGTTTCTACCTTCGCGGTTTTACTTGAGTGACGCTGCTCATGCTTGCAA

CCGCTGGGATGCAGGTGCATGCCTCTAGCATGAAGTCAGACAA

T(InsC2)

**Supplementary Figure S3:** ITS2 variants detected in coral samples. ITS2 variants of D1, C40, C66 and C15, are indicated with a specific colour and their corresponding ITS2 sequence name. Ins= nucleotide insertion, *= nucleotide deletion.
